# The effect of population bottleneck size and selective regime on genetic diversity and evolvability in bacteria

**DOI:** 10.1101/726158

**Authors:** Tanita Wein, Tal Dagan

## Abstract

Population bottlenecks leading to a drastic reduction of the population size are common in the evolutionary dynamics of natural populations; their occurrence is known to have implications for genome evolution due to genetic drift, the consequent reduction in genetic diversity and the rate of adaptation. Nevertheless, an empirical characterization of the effect of population bottleneck size on evolutionary dynamics of bacteria is currently lacking. Here we show that selective conditions have a stronger effect on the evolutionary history of bacteria in comparison to genetic drift following population bottlenecks. We evolved *Escherichia coli* populations under three different population bottlenecks (small, medium, large) in two temperature regimes (37°C and 20°C). We find a high genetic diversity in the large in comparison to the small bottleneck size. Nonetheless, the cold temperature led to reduced genetic diversity in all bottleneck sizes, hence, the temperature has a stronger effect on the genetic diversity in comparison to the bottleneck size. A comparison of the fitness gain among the evolved populations reveals a similar pattern where the temperature has a significant effect on the fitness. Our study demonstrates that population bottlenecks are an important determinant of the evolvability in bacteria; their consequences depend on the selective conditions and are best understood via their effect on the standing genetic variation.

## Introduction

Population bottlenecks impose a rapid and often drastic reduction on the population size. Such events are common and unavoidable during the evolutionary dynamics of natural populations. Fluctuations in the size of the population over time play a major role in the ecology and evolution of prokaryotic organisms (Fraser et al. 2009). Abiotic factors in the environment can lead to severe population size reduction. Examples are seasonal change (e.g. temperature fluctuations) or resource limitation that can result in only a small fraction of a population surviving the temporary selective event (e.g. via persistence or dormancy (Balaban et al. 2004)). Species interactions (i.e., biotic factors) may also lead to fluctuations in bacterial population size over time. For example, the life cycle of host-associated bacteria is often characterized by repeated population bottleneck events. For pathogenic bacteria, the transmission to a new host and the selection by the host immune system drastically reduces bacterial population size in every infection cycle (Didelot et al. 2016; Moxon and Kussell 2017). The life cycle of bacteria in mutualistic interactions (i.e., beneficial symbiosis) is also characterized by successive population bottlenecks, which typically occur at the initial stages of the host colonization due to founder effects. The effect of strong population bottlenecks is well recognized in vertically inherited bacterial symbionts, where only few bacterial cells are transferred to the next generation (e.g., as in aphid or beetle symbioses (Funk et al. 2001; McCutcheon and Moran 2011; Salem et al. 2017). The evolution of horizontally transmitted symbionts can be characterized by strong population bottlenecks as well. Next to founder effects, priority effects in colonization may induce strong population bottlenecks for successively incoming colonizers, where the first colonizer restricts the habitat for the later incoming colonizers (Stephens et al. 2015; Wein et al. 2018). Finally, phage predation constitutes a major factor leading to repeated population bottlenecks and consequently fluctuating populations size of bacterial populations (Avrani et al. 2011; Koskella and Brockhurst 2014). Notably, population bottlenecks may be either neutral due to random sampling of the bacterial population, or selective – where the probability of surviving the bottleneck depends on the genotype.

The occurrence of population bottlenecks is known to have significant implications for bacterial genome evolution due to their potential to lead to genetic drift, which results in a reduction in the population genetic diversity. This is true for both types of bottlenecks, where in the adaptive bottleneck the effect of drift is limited to alleles in linkage with the selected genotype. The stochastic elimination of rare alleles from the population during genetic drift may also have consequences for the rate of adaption as they decrease the efficacy of natural selection (Brandvain and Wright 2016). Strong (or small) population bottlenecks are assumed to maintain slightly deleterious mutations in the population (Lynch et al. 1993) and thus decrease the rate of adaptation, while weak (or large) population bottlenecks are expected to maintain a higher rate of adaptation. However, in large populations that experience weak neutral population bottlenecks, independently derived beneficial mutations are expected to compete in the population. The relative frequency of such beneficial mutations in the population has a direct effect on the dynamics of less beneficial mutations due to linkage disequilibrium. Under adaptive bottlenecks (i.e., strong selective conditions), this phenomenon – which has been termed Hill–Robertson effect, or clonal interference (Gerrish and Lenski 1998; Hill and Robertson 2009) – can thus lead to bacterial adaptation that is driven by highly beneficial mutations and is characterized by a high probability of parallel evolution (Herron and Doebeli 2013). The dynamics of clonal interference may be nonetheless perturbed in the presence of population bottlenecks where strong bottlenecks are expected, in addition, to decrease the probability of parallel evolution (Wahl et al. 2002).

Population bottlenecks thus constitute an important determinant of allele dynamics that interferes with selective processes, including purifying selection of deleterious alleles as well as positive selection for beneficial alleles. Nonetheless, a quantification of the combined effect of population bottlenecks and selection regimes on allele frequency dynamics remains challenging due to the multiplicity of confounding factors. Here we compare properties of genome and phenotype evolution in *Escherichia coli* under three different population bottleneck sizes and two different selection regimes. Theory predicts that the population size, as imposed by the strength of the bottleneck events, has an effect on the genetic diversity depending on the environmental conditions (i.e., selection regime (Lanfear et al. 2014)). The level of adaptation is similarly expected to vary among the different combinations of population bottleneck size and selection regimes; the highest increase in fitness (i.e., rapid adaptation) is expected in the largest population bottleneck size and harshest selective conditions.

## Results

To test the theoretical predictions, we conducted an experimental evolution study of *E. coli* strain K-12 MG1655 in a serial transfer approach. The three bottleneck sizes were applied in every serial transfer with a dilution of 1:100 (1% of the total population, large, L), 1:1,000 (0.1% of the total population, medium, M) and 1:10,000 (0.01% of the total population, small, S). The populations were evolved in two environmental temperatures: 37°C that is considered as optimal growth conditions and 20°C that is considered sub-optimal conditions for *E. coli*. The experiment was conducted with eight replicates for approximately 1,000 generations. Comparative genomics of the evolved and ancestral populations revealed 208 evolved single nucleotide variants (SNVs; using a threshold of SNV allele frequency (AF) ≥0.02), of which 81 (39%) SNVs were observed in more than one population (Table S1).

### High genetic diversity in the large bottleneck and reduction of diversity at 20°C

To examine the effect of bottleneck size and temperature on the degree of genetic polymorphism we compared the nucleotide diversity among the evolved populations. Our results revealed a significant effect of the temperature and bottleneck size on the population nucleotide diversity (!1A). Evolution in 20°C resulted in overall low genetic diversity in all populations compared to populations evolved at 37°C. The highest genetic diversity was observed in L populations and the lowest was observed in M populations in both temperature regimes (Fig. 1A). Furthermore, our results show that there is no interaction between the effects of bottleneck size and temperature on the mean population nucleotide diversity; hence, the effect of the bottleneck was similar in both temperature regimes (Fig. 1A). Consequently, we conclude that the population bottleneck size had an impact on the genetic diversity regardless the selection pressure imposed by the temperature in our experiment.

**Figure 1.**
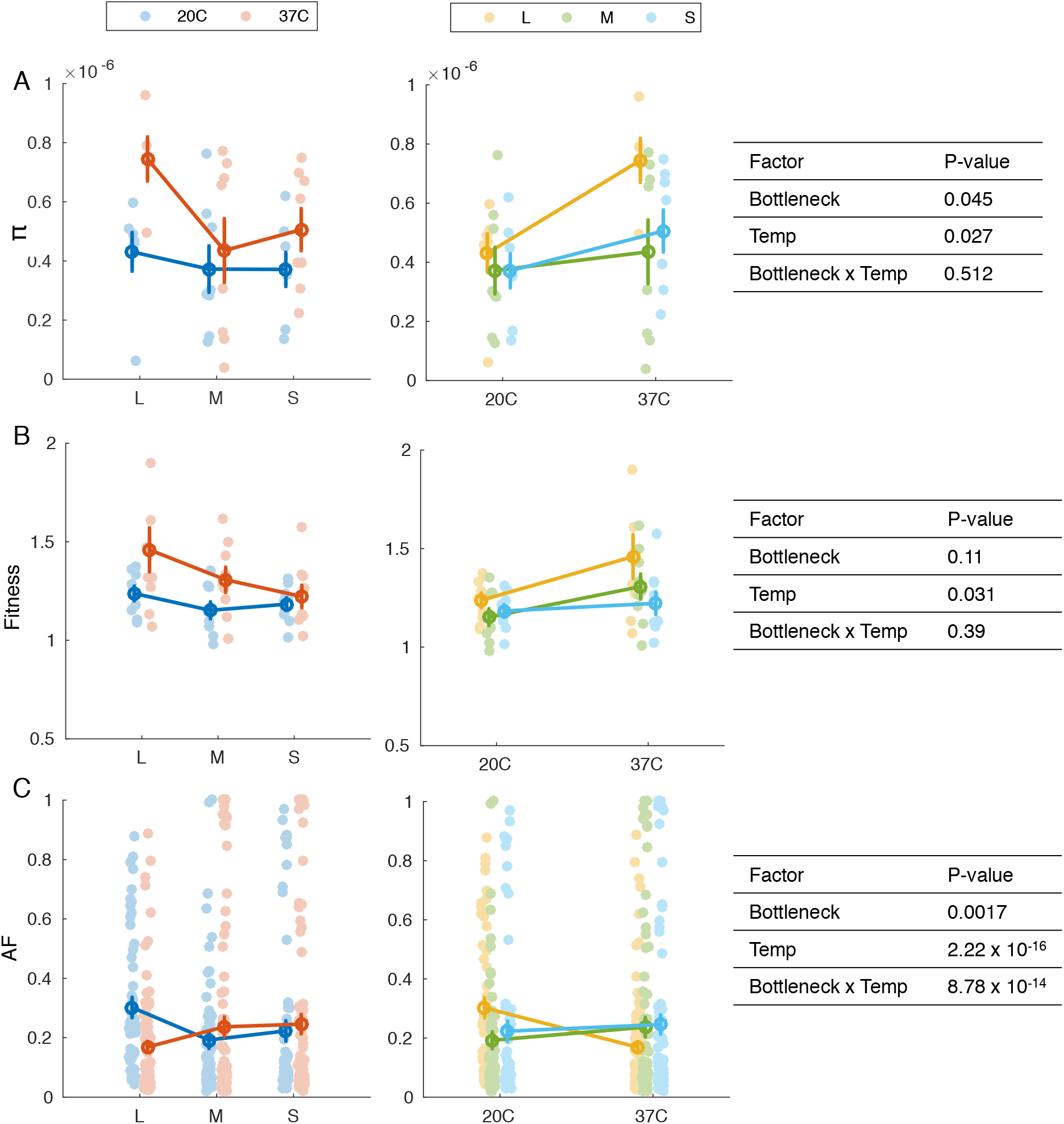
The effect of bottleneck size and temperature on (A) nucleotide diversity (π) (B) relative fitness and (C) allele frequency (AF). Bottleneck size is denoted as L: Large, M: Medium, or S: Small. The data is presented by dots; mean values ± SEM are marked by a circle with error bars. Note that π and fitness are presented per population, while AF is presented for all mutated loci in all populations. Tables on the right show the results of ANOVA two-way performed on transformed data (using aligned transformation (Wobbrock et al. 2011)). The quality of the transformation was validated by ANOVA; All transformations yielded F-values close to 0 as required. The statistical test of the effect on AF was performed including only parallel loci (i.e., evolved in ≥ 1 population) and the populations were considered as replicates. Fitness values >1 indicate a relative fitness increase of the evolved population compared to the ancestral population.

### Significant effect of temperature on the fitness of evolved populations

To test the effect of bottleneck size on adaptive evolution in our experiment, we compared the fitness of the evolved populations relative to the ancestral population. For that purpose, we conducted competition experiments between a marked ancestor and the evolved populations (in the corresponding temperature regime). This revealed that the tested populations had an increased fitness relative to the ancestral population (Fig. 1B). Our results further show that the effect of temperature on the evolved populations fitness is significant. The relative fitness increase of the populations evolved at 37°C was higher in comparison to the populations evolved at 20°C (Fig. 1B). In contrast, the bottleneck size had not significant effect on the evolved populations relative fitness (Fig. 1B; but we note that a trend of increased fitness with population size bottleneck at 37°C can be observed). Furthermore, there is no significant interaction between the effects of bottleneck size and temperature on the mean relative fitness; hence, the effect of bottleneck size was similar in both temperature regimes (Fig. 1B). Notably, the effect of bottleneck size and temperature on the relative fitness was similar to what we observed in the comparison of genetic diversity among the evolved populations (Fig. 1A).

### Allele frequency dynamics depend on both bottleneck size and temperature

Comparing the allele frequency of SNVs in the evolved populations we found that both temperature and population bottleneck size had an effect on the distribution of SNV allele frequency in the population (Fig. 1C). Furthermore, the effect of bottleneck size on the allele frequency (AF) varied among the growth temperatures. Indeed, we found a significant interaction between temperature and bottleneck size; hence, the effect of bottleneck size on the allele frequency depends on the growth temperature (and *vice versa*). While the AF distribution in M and S populations was similar in both temperature regimes, the L populations evolved at 37°C (L37) where characterized by lower AF in comparison to L populations evolved at 20°C (L20; Fig. 1C). A comparison of AF distribution among synonymous and non-synonymous SNVs further showed that the observed differences in AFs are well explained by the allele dynamics of non-synonymous rather than synonymous SNVs (Fig. S1). Notably, no SNVs reached fixation (i.e., AF>0.9) in the L37 populations, while several SNVs reached a high frequency (AF>0.9) in the M and S populations at both growth temperatures (Fig. 1C; Table S1).

The fixed mutations include several genetic variants that may be linked to osmotic stress (triggered by the salt concentration). Examples are fixed mutations observed in the Sodium/glutamate symporter gene (*gltS*; M37) and the potassium uptake system. These include mutations in *trkH* or *trkD* (*kup*) that were fixed across the three bottleneck sizes at 37°C (but not all replicates), as well as mutations in *sapD*, which were fixed only in the small bottleneck size of both temperature regimes. Further fixed mutations targeting transcriptional processes occurred across temperatures and bottleneck sizes in the DNA-dependent RNA polymerase gene (*rpoC*), while fixed mutations in *rpoB* only occurred in L20 populations. Mutations related to translational processes occurred only at 20°C in genes encoding for ribosomal proteins, *rpsG* and *rpsA*. In addition, fixed mutations emerged in genes involved in nutrient abundance and starvation response (e.g., *spoT* and *sspA* in L20 populations) or regulation of anoxic metabolism (e.g. *arcA* in one M37 population). All of the fixed variants described above are non-synonymous substitutions.

To further examine differences in AF dynamics depending on the evolutionary factors we examined the fate of preexisting SNVs in the ancestral population. The ancestral *E. coli* strain in our population had four SNVs in comparison to the reference genome; of which two were at a high AF: a synonymous substitution in *aroE* (AF_ancestral_=0.98) and a synonymous substitution in *yciM* (AF_ancestral_=0.77) (Table S1). These two substitution reached fixation (AF>0.9) in all evolved populations (Fig. S2). One intergenic mutation remained either at a similar low allele frequency (AF≈0.3) in some populations or slightly decreased in others regardless of the conditions or bottleneck size. This intergenic ancestral variant is thus likely to be selectively neutral. An additional ancestral synonymous SNV in *rpoD* that was present with an AF=0.2 decreased in frequency in almost all populations. RpoD is sigma factor that is involved in translation during exponential growth (e.g. ribosomal operons or rRNA- and tRNA genes). The allele dynamics of the RpoD SNV indicate that this variant evolved under purifying selection.

### Large populations bottlenecks show the highest degree of parallel evolution

The distribution of shared SNVs among replicate populations evolved under the same evolutionary factors revealed that more parallel SNVs were found among replicates in the L populations in comparison to the M and S populations at both temperature regimes (Fig. S3). The parallel SNVs in L populations were, however, specific to the temperature regime; this is true for the specific mutation position as well as for the gene in which the mutation occurred (Fig. 2; Fig. S3). Notably, the comparison among populations evolved under the same temperature did not reveal a strong signal of parallel evolution. Nevertheless, at 37°C we observed more parallel variants across bottleneck sizes than at 20°C (Fig. S3). Moreover, several of the variants detected as parallel in a specific bottleneck, e.g., S20 or S37, did not occur in another bottleneck size and can thus be considered bottleneck size specific (Fig. 2).

**Figure 2.**
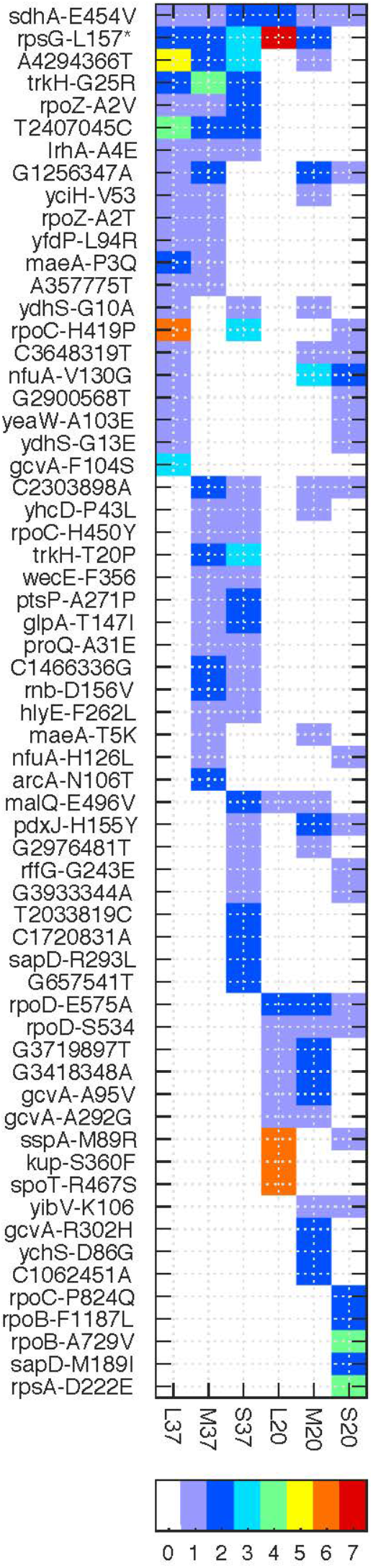
The distribution and annotation of parallel SNVs in the evolved populations. SNVs that have been observed in >1 replicate population (AF≥0.02) are presented with the number of replicated color coded according to the colorbar at the bottom. The variant annotation is listed on the left according to the following format: intergenic variants are presented by [ancestral nucleotide][genomic position][evolved nucleotide]. Intragenic variants are presented with the gene symbol [ancestral amino acid][amino acid position][evolved amino acid]. For synonymous SVNs, the evolved amino acid is omitted; STOP codons are marked by ‘*’. Ancestral variants are excluded.

One variant was detected across all bottleneck sizes and both temperatures. The nonsynonymous SNV in *sdhA* was observed in eight replicate populations across all conditions (*sdhA*-E454V). SdhA is involved in the synthesis of fumarate from succinate and can switch function between aerobic and anaerobic metabolism (Ruprecht et al. 2009). Several SNVs are specific to the temperature regime (yet not across all bottleneck sizes or replicates). For example, at 37°C, this includes variants in genes related to transcriptional processes such as the RNA-polymerase subunit gene *rpoZ* as well as the transcriptional regulator *lrhA*. At 20°C, we observed two more variants in *rpoD* that did not emerge at 37°C. Overall many parallel occurring variants include genes associated with growth (e.g. *rpoB, rpoC, rpoZ*) as well as persistence in the extended stationary growth phase such as the anoxic regulation protein system *arcA/B* or proteins involved in starvation response, e.g. *spoT* encoding for bifunctional (p)ppGpp synthase/hydrolase that is activated in response to nutritional changes and *sspA* encoding for the stringent starvation protein. Notably, only at 20°C we observed fixed parallel substitutions in the ribosomal protein RpsG that is involved in mRNA binding (i.e., translation) which were shared across the M and L bottleneck size indicating its relation to the temperature regime. Overall, most of the parallel SNVs were observed in the L populations regardless of the temperature, yet, parallel SNVs across bottlenecks were more frequent in populations evolved at 37°C.

## Discussion

The evolutionary trajectory of populations through time is influenced by the interplay of drift and selection that act on the standing genetic variation. In the context of sexually reproducing eukaryotes, the recognition that population bottlenecks reduce genetic variation was made long ago (e.g., (Mayr 1963)) while Nei et al. (1975) quantified the extent of loss of variation. Our results are in agreement with those observations, and extend their applicability to the prokaryotic domain, since populations evolved under a large bottleneck maintain the highest genetic diversity over time. Nonetheless, our results show that the differences between the medium or small bottleneck are only marginal. The lack of difference between those bottlenecks suggests that the small population bottleneck in our experiment (10^5^ cells) already includes a sufficient number of variants for maintaining a basic level of standing genetic variation and by that avoid a great loss of diversity within the population. Notably, the number of fixed mutations (i.e., AF≥0.9) is highest in the S populations and lowest in the L populations (Fig. 1C). The difference in the frequency of fixed SNVs may be explained by a strong effect of competing beneficial mutations in the L populations (i.e., clonal interference), and in addition, a strong impact of genetic drift in the S populations (Fig. 1C).

The degree of standing genetic variation is expected to impose a fundamental constraint on the rate of adaptation. Our results indicate that both genetic variation and adaptation (i.e., fitness) are highest in the large bottleneck. In addition, we observe a higher number of non-synonymous mutations at 37°C, that is in line with the higher increase in fitness in the evolved populations at 37°C but not at 20°C. In the cold temperature regime some highly adaptive mutations may induce selective sweeps reducing the diversity and increasing the number of fixed mutations. Our results thus show that the genetic diversity, the frequency of non-synonymous mutations and fitness are tightly linked and may be of assistance when predicting the degree of adaptation in other (natural) settings.

Additionally, we observed that the degree of parallel evolution is sensitive to the bottleneck size regardless of the environment, contrasting previous studies on serial bottlenecking (e.g., (Vogwill et al. 2016)). Theory predicts that the degree of parallel evolution is expected to be highest in large populations compared to small populations (Orr 2005). Thus, our findings for the effect of bottleneck size on parallel evolution are in agreement with theoretical predictions for the effect of constant populations size.

The genes evolved in our experiment reveal a strong impact of our experimental system. The high abundance of variants related to fast growth (e.g, *rpo* genes) in the evolved populations indicates that the exponential growth phase, which is typical for serial transfer experiments, imposes a strong selection pressure on traits related to fast growth regardless of the temperature or bottleneck size. In addition, genes related to persistence and nutrient starvation are important in the stationary growth phase of bacteria and thus may be important for growth in the extended stationary phase, especially in the L populations. Mutations in genes related to such adaptation seem to evolve under strong positive selection and their allele dynamics may mask other low frequency variants that emerged during the experiment. We propose that the serial transfer approach is likely to select for the mutations that reached the highest frequencies (i.e., of fast growers) and thus reduce the overall genetic diversity. Thus, upon the induction of the bottleneck effect (i.e., the transfer), high frequency variants will increase while low frequency variants may quickly disappear. This implies that the probability of fixation to occur in the population is not uniformly distributed across growth phases; mutations that emerge early in the growth phase have a higher probability of being fixed, while mutations that emerge at a later stage are a minority and therefore less likely to be fixed. This suggestion is in agreement with previously published mathematical models of the survival probability of mutations in bacterial populations grown in batch cultures (Wahl and Zhu 2015). We conclude that the differences in the growth dynamics (i.e., mutational dynamics along growth phases) between the large, and medium/small populations may have a significant effect on the allele dynamics in the population, even to a larger extent than the genetic drift introduced by the serial population bottlenecks.

Our results indicate that selection remains a major force of evolutionary dynamics and selective conditions may have a stronger effect on the evolutionary history of bacteria in comparison to repeated bottlenecks of various sizes. Nevertheless, serial bottleneck effects remain an important determinant in the evolution of bacterial populations, especially with regards to the rate of adaptation. An example for the importance of population bottlenecks in natural habitats is the impact of phage predation on the evolution of their bacterial hosts. It is well known that phage predation imposes a strong population bottleneck on the host and several studies showed that this may lead to rapid evolution of phage resistance (Paterson et al. 2010). Nonetheless, our results suggest that while the strong population bottleneck will lead to the fixation of specific genotypes (i.e., the resistant genotypes; as in our S bottleneck (Fig 1C)), at the same time, the bottleneck will lead to a purge of the host genetic diversity (as in the S bottleneck, Fig. 1A). Taken together, we expect that the adaptability of the host population to other selection pressures (e.g., abiotic factors in the environment) will be significantly reduced due to the repeated bottlenecks. Indeed, previous studies observed that the adaptability of bacterial populations is reduced when exposed to multiple stressors (e.g., phage and fast growth or predation and antibiotics (Avrani and Lindell 2015; Hiltunen et al. 2018)). Similarly, selection events for the dissemination of plasmids encoding for antibiotics (or metal) resistance may also impose a strong population bottleneck on the plasmid carrying cells. Fluctuating selective conditions for the plasmid-encoded function (i.e., antibiotics) are parallel to serial population bottlenecks for the portion of the population that carries the plasmid (Wein et al. 2019). Also here, the strong selection for the plasmid presence may lead to the fixation of successful plasmid-host genotypes (e.g. (De Gelder et al. 2008; Harrison et al. 2015)), but, at the same time, to a reduction in the population genetic diversity, which is expected to decrease the rate of adaptation to alternative selection pressures in the environment. Thus, population bottlenecks induced by abiotic and biotic factors are expected to have a significant effect on the adaptability of bacterial population and their consequences are best understood via their effect on the standing genetic variation.

## Materials and Methods

### Experimental evolution

The experiment was conducted with the *E. coli* K-12 strain MG1655. Eight ancestral replicates were picked as single colonies from LB-agar plates and inoculates for overnight growth at 37°C. Thereafter, the ancestral cultures were divided into two temperatures of 37°C and 20°C and three bottlenecks sizes. The population bottleneck sizes were applied every serial transfer with a dilution of 1:100 (1%, large, L), 1:1,000 (0.1%, medium, M) and 1:10,000 (0.01%, small, S). The replicated populations were evolved in a total volume of 1 ml LB medium. The populations were propagated either every 12 h at 37°C or every 24 h at 20°C according to their growth dynamics for a total of 130 for L, 110 for M and 90 for S transfers (i.e., bottlenecks) and approximately 1000 generations.

### Sequence analysis

Population sequencing was applied to enable the detection of variant alleles. Total DNA was isolated from 1 ml culture using the Wizard Genomic DNA Purification Kit (Promega). Concentration and quality of the extracted DNA was assessed using the NanoDrop™ (Thermo Fisher Scientific) and Qubit (Invitrogen by Life Technologies). The sample libraries for Illumina sequencing were prepared using the Nextera DNA Flex and the Nextera XT library kit (Illumina, Inc) and sequencing was performed with paired end reads on the Hiseq system (Illumina, Inc). The median coverage in all populations ranged between x55 and x255.

Sequencing reads were trimmed to remove Illumina specific adaptors and low quality bases using the program Trimmomatic v.0.35 (Bolger et al. 2014) (parameters: NexteraPE-PE.fa:2:30:10 CROP:125 HEADCROP:5 LEADING:5 TRAILING:5 SLIDINGWINDOW:4:20 MINLEN:36). The sequencing reads were mapped to the reference genomes using BWA-MEM 0.7.16a-r1181 (Li and Durbin 2009). As the reference we used the *E. coli* MG1655 genome (GenBank acc. no. NC_000913.3). Mapping statistics were retrieved using Alfred v0.1.5 (Rausch et al. 2018) Subsequent indexing and local realignment of sequencing reads were performed using PICARD tools, SAMtools v1.6 (Li et al. 2009) and GATK v3.8-0-ge9d806836 (McKenna et al. 2010). Short indels and SNPs were called using LoFreq v.2.1.2 (Wilm et al. 2012). The annotation of evolved SNVs was performed with an in-house PERL script.

The analysis of the evolved population revealed cross contamination in 4 populations that were excluded from further analysis. Data analysis and statistical tests were performed with MatLab© and R version 3.5.1. Data transformation for the statistical tests was performed with the ARTtool package (Wobbrock et al. 2011).

Nucleotide diversity (π) was calculated as in (Schloissnig et al. 2013): 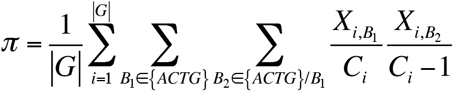, where *G* is the length of *E. coli* reference genome, and *X*_*i,Bj*_ is the count of a specific nucleotide *B*_*j*_ at a specific locus *i* with coverage *C*_*i*_.

### Competition experiments

The relative fitness (*w*, (Lenski et al. 1991)) of the evolved versus the ancestral strain (marked) was estimated by direct pairwise competition experiments, with three replicates per three populations per bottleneck and temperature. The impact of the marked *E. coli* MG1655 Tm^r^ strain was previously evaluated by competition experiments of the wildtype against the marked strain (Wein et al. 2019). All competition experiments were initiated with a 1:1 mixture of 1:100 diluted evolved strain and ancestral strain from overnight cultures, in a total volume of 1 ml of LB medium. The relative fitness of the evolved strains was calculated by gaining viable cell counts at the time points 0 h, 24 h and 48 h. The strains were distinguished through plating on non-selective (LB) and selective media (LB supplemented with trimethoprim 150 µg/ml).

## Data availability

Sequence reads available in SRA Accessions: ####.

## Supplemental material

Figures S1, S2, S3, Table S1.

## Acknowledgments

With thank Giddy Landan, Anne Kupczok, Elie Jami, Maxime Godfroid, Ana Garona and Nils Hülter for fruitful discussions and critical comments on the manuscript. Genome sequencing was performed in the Centre for Genome Analysis Kiel that is funded by the German Research Foundation (DFG). The study was supported by the ZMB Young Scientist Grant 2018 (awarded to T.W.) and the DFG focus program 1819 (Grant no. DA1202/2-1; awarded to T.D.).

## References

Avrani S, Lindell D. 2015. Convergent evolution toward an improved growth rate and a reduced resistance range in *Prochlorococcus* strains resistant to phage. Proc. Natl. Acad. Sci. USA 112:E2191–E2200.

Avrani S, Wurtzel O, Sharon I, Sorek R, Lindell D. 2011. Genomic island variability facilitates *Prochlorococcus–*virus coexistence. Nature 474:604–608.

Balaban NQ, Merrin J, Chait R, Kowalik L, Leibler S. 2004. Bacterial persistence as a phenotypic switch. Science 305:1622–1625.

Bolger AM, Lohse M, Usadel B. 2014. Trimmomatic: a flexible trimmer for Illumina sequence data. Bioinformatics 30:2114–2120.

Brandvain Y, Wright SI. 2016. The limits of natural selection in a nonequilibrium world. Trends in Genetics 32:201–210.

De Gelder L, Williams JJ, Ponciano JEM, Sota M, Top EM. 2008. Adaptive plasmid evolution results in host-range expansion of a broad-host-range plasmid. Genetics 178:2179–2190.

Didelot X, Walker AS, Peto TE, Crook DW, Wilson DJ. 2016. Within-host evolution of bacterial pathogens. Nat. Rev. Micro. 14:150–162.

Fraser C, Alm EJ, Polz MF, Spratt BG, Hanage WP. 2009. The bacterial species challenge: making sense of genetic and ecological diversity. Science 323:741–746.

Funk DJ, Wernegreen JJ, Moran NA. 2001. Intraspecific variation in symbiont genomes: bottlenecks and the aphid-buchnera association. Genetics 157:477–489.

Gerrish PJ, Lenski RE. 1998. The fate of competing beneficial mutations in an asexual population. Genetica 102-103:127–144.

Harrison E, Guymer D, Spiers AJ, Paterson S, Brockhurst MA. 2015. Parallel compensatory evolution stabilizes plasmids across the parasitism-mutualism continuum. Curr. Biol. 25:2034–2039.

Herron MD, Doebeli M. 2013. Parallel evolutionary dynamics of adaptive diversification in *Escherichia coli*. PLoS biol. 11:e1001490.

Hill WG, Robertson A. 2009. The effect of linkage on limits to artificial selection. Genet. Res. 8:269–294.

Hiltunen T, Cairns J, Frickel J, Jalasvuori M, Laakso J, Kaitala V, Künzel S, Karakoc E, Becks L. 2018. Dual-stressor selection alters eco-evolutionary dynamics in experimental communities. Nat. ecol. evol. 2:1974–1981.

Koskella B, Brockhurst MA. 2014. Bacteria-phage coevolution as a driver of ecological and evolutionary processes in microbial communities. FEMS Microbiol. Rev. 38:916–931.

Lanfear R, Kokko H, Eyre-Walker A. 2014. Population size and the rate of evolution. Trends Ecol. Evol. 29:33–41.

Lenski RE, Rose MR, Simpson SC, Tadler SC. 1991. Long-term experimental evolution in *Escherichia coli*. I. Adaptation and divergence during 2,000 generations. Amer. Nat. 138:1315–1341.

Li H, Durbin R. 2009. Fast and accurate short read alignment with Burrows-Wheeler transform. Bioinformatics 25:1754–1760.

Li H, Handsaker B, Wysoker A, Fennell T, Ruan J, Homer N, Marth G, Abecasis G, Durbin R, 1000 Genome Project Data Processing Subgroup. 2009. The sequence alignment/map format and SAMtools. Bioinformatics 25:2078–2079.

Lynch M, Bürger R, Butcher D, Gabriel W. 1993. The mutational meltdown in asexual populations. J. Hered. 84:339–344.

Mayr E. 1963. Animal Species and Evolution. Harvard University Press 1362–4962.

McCutcheon JP, Moran NA. 2011. Extreme genome reduction in symbiotic bacteria. Nat. Rev. Micro. 10:13–26.

McKenna A, Hanna M, Banks E, Sivachenko A, Cibulskis K, Kernytsky A, Garimella K, Altshuler D, Gabriel S, Daly M, et al. 2010. The genome analysis toolkit: A MapReduce framework for analyzing next-generation DNA sequencing data. Genome research 20:1297–1303.

Moxon R, Kussell E. 2017. The impact of bottlenecks on microbial survival, adaptation, and phenotypic switching in host-pathogen interactions. Evolution 71:2803–2816.

Nei M, Maruyama T, Chakraborty R. 1975. The bottleneck effect and genetic variability in populations. Evolution 29:1.

Orr HA. 2005. The probability of parallel evolution. Evolution 59:216–220.

Paterson S, Vogwill T, Buckling A, Benmayor R, Spiers AJ, Thomson NR, Quail M, Smith F, Walker D, Libberton B, et al. 2010. Antagonistic coevolution accelerates molecular evolution. Nature 464:275–278.

Rausch T, Fritz MH-Y, Korbel JO, Benes V. 2018. Alfred: Interactive multi-sample BAM alignment statistics, feature counting and feature annotation for long- and short-read sequencing. Bioinformatics 28:1530.

Ruprecht J, Yankovskaya V, Maklashina E, Iwata S, Cecchini G. 2009. Structure of *Escherichia coli* succinate: quinone oxidoreductase with an occupied and empty quinone-binding site. J. Biol. Chem. 284:29836–29846.

Salem H, Bauer E, Kirsch R, Berasategui A, Cripps M, Weiss B, Koga R, Fukumori K, Vogel H, Fukatsu T, et al. 2017. Drastic genome reduction in an herbivore’s pectinolytic symbiont. Cell 171:1–12.

Schloissnig S, Arumugam M, Sunagawa S, Mitreva M, Tap J, Zhu A, Waller A, Mende DR, Kultima JR, Martin J, et al. 2013. Genomic variation landscape of the human gut microbiome. Nature 493:45–50.

Stephens WZ, Wiles TJ, Martinez ES, Jemielita M, Burns AR, Parthasarathy R, Bohannan BJM, Guillemin K. 2015. Identification of population bottlenecks and colonization factors during assembly of bacterial communities within the zebrafish Intestine. mBio 6:e01163- 15–11.

Vogwill T, Phillips RL, Gifford DR, Maclean RC. 2016. Divergent evolution peaks under intermediate population bottlenecks during bacterial experimental evolution. Proc. R. Soc. B 283:20160749.

Wahl LM, Gerrish PJ, Saika-Voivod I. 2002. Evaluating the impact of population bottlenecks in experimental evolution. Genetics 162:961–971.

Wahl LM, Zhu AD. 2015. Survival probability of beneficial mutations in bacterial batch culture. Genetics 200:309–320.

Wein T, Dagan T, Fraune S, Bosch TCG, Reusch TBH, Hülter NF. 2018. Carrying capacity and colonization dynamics of *Curvibacter* in the *Hydra* host habitat. Front. Microbiol. 9:443.

Wein T, Hülter NF, Mizrahi I, Dagan T. 2019. Emergence of plasmid stability under non-selective conditions maintains antibiotic resistance. Nat. Commun. 10:2595.

Wilm A, Aw PPK, Bertrand D, Yeo GHT, Ong SH, Wong CH, Khor CC, Petric R, Hibberd ML, Nagarajan N. 2012. LoFreq: a sequence-quality aware, ultra-sensitive variant caller for uncovering cell-population heterogeneity from high-throughput sequencing datasets. Nuc. Ac. Res. 40:11189–11201.

Wobbrock JO, Findlater L, Gergle D, Higgens JJ. 2011. The aligned rank transform for nonparametric factorial analyses using only ANOVA procedures. New York ACM:143–146.

