## Supplementary figures for "The effect of population bottleneck size and selective regime on genetic diversity and evolvability in bacteria"

**Figure S1.** The effect of bottleneck size and temperature on allele frequency (AF) of intergenic, synonymous, and nonsynonymous SNVs. Bottleneck size is denoted as L: Large, M: Medium, or S: Small. The data is presented by dots; mean values ± SEM are marked by a circle with error bars.

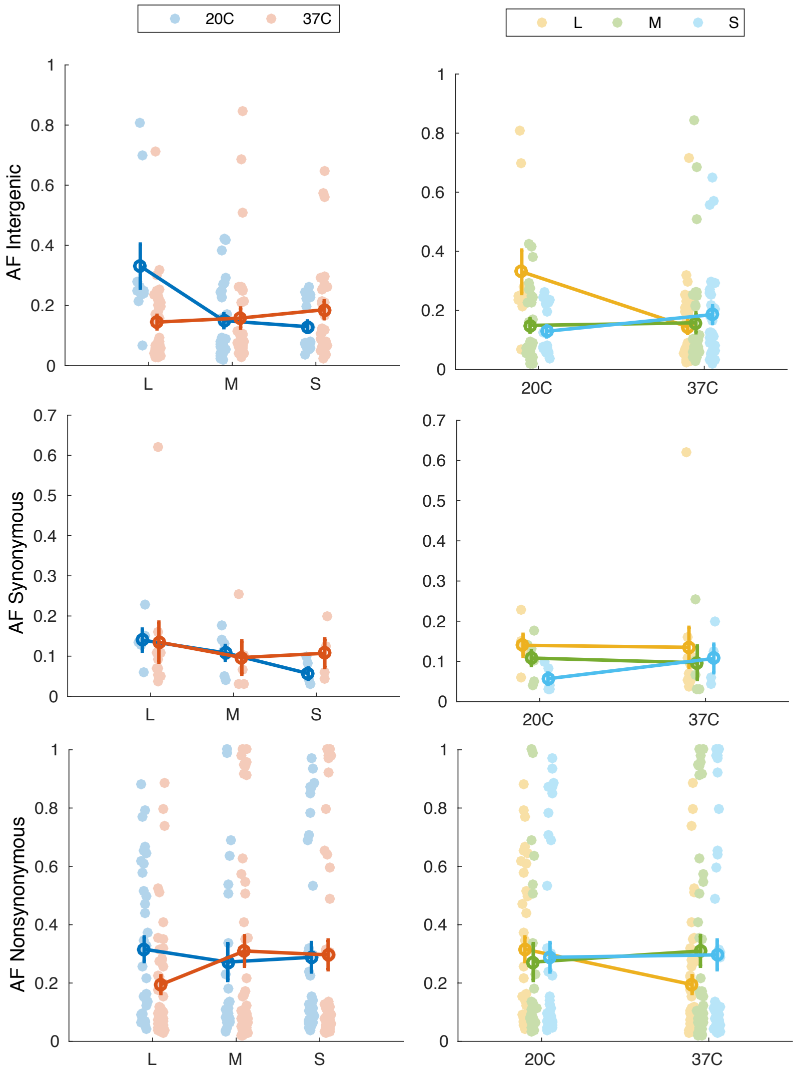

**Figure S2.** Distribution of ancestral variant allele frequency in the evolved populations. The ancestral allele frequency is marked by a green line. The distribution of allele frequency in the evolve populations is shown by dots.

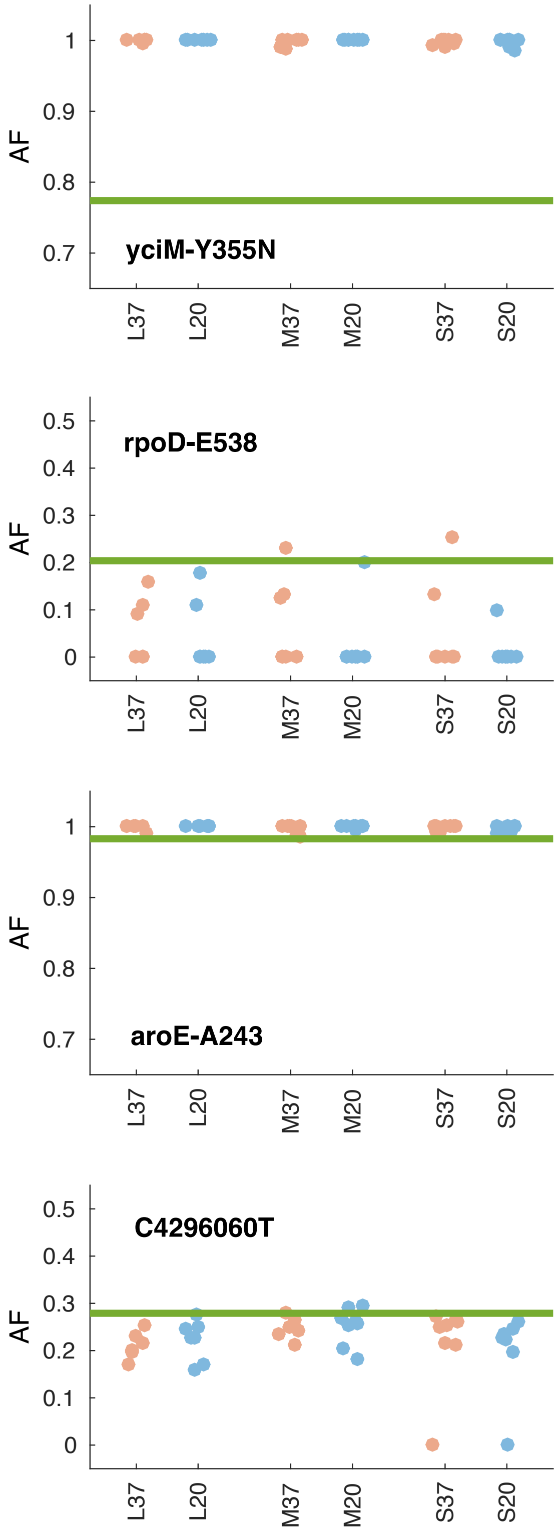

**Figure S3**. The distribution of shared SNV loci among replicate populations. Cell color is the number of shared mutations (see colorbar at the bottom).

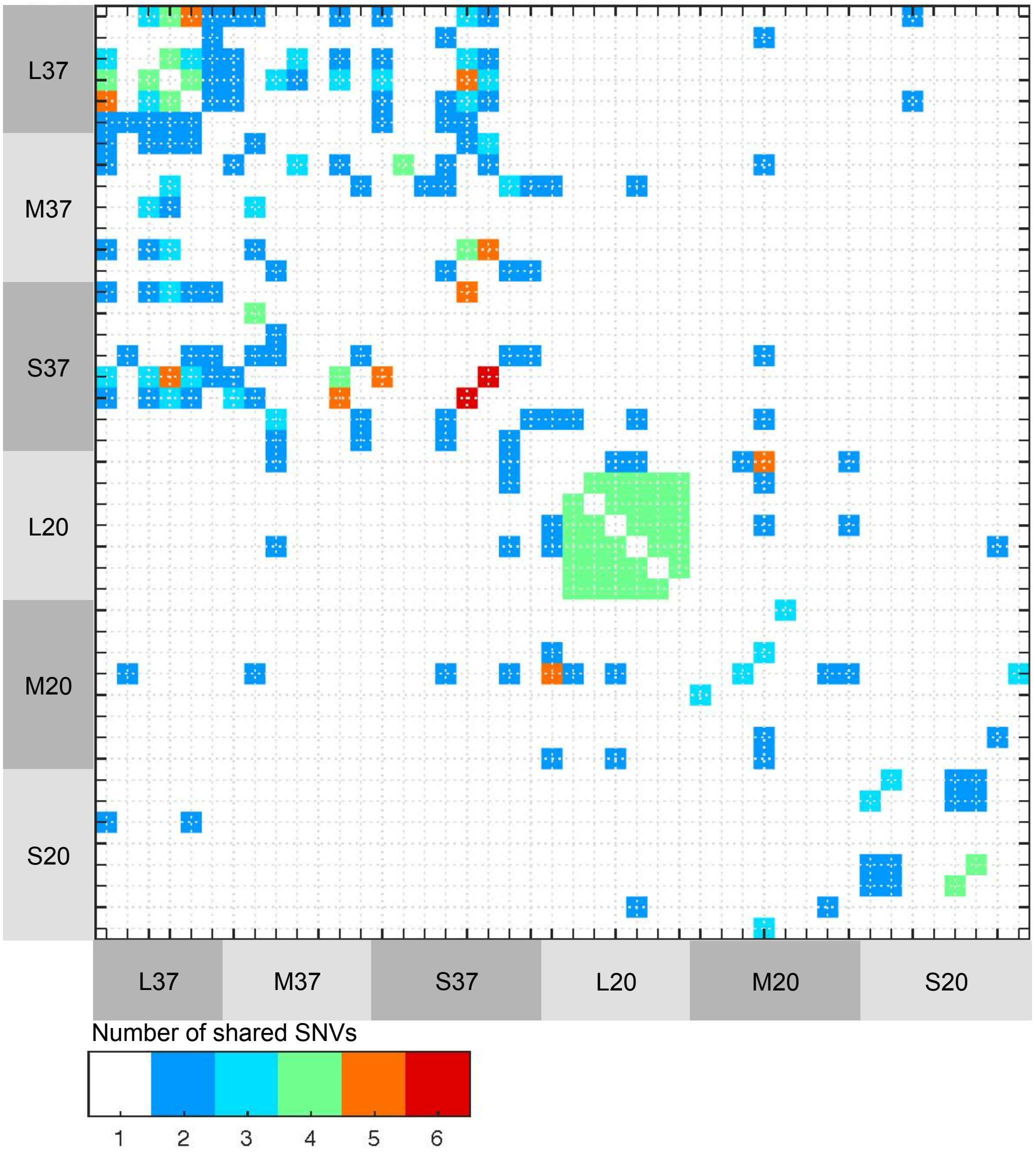
